# Differential effects of prolonged aerobic and resistance exercise on cognitive functioning in sedentary young adults

**DOI:** 10.1101/2023.04.03.535439

**Authors:** Francesco Riganello, Alexandra Pearce, Kathleen M. Lyons, Adrian M. Owen, Andrea Soddu, Bobby Stojanoski

**Affiliations:** S. Anna Institute and Research in Advanced Neurorehabilitation (RAN), Crotone, Italy; Anesthesiology Institute, Cleveland Clinic, Cleveland, Ohio; Western Institute for Neuroscience, Department of Physiology and Pharmacology and Department of Psychology, University of Western Ontario, London ON, Canada; Western Institute for Neuroscience, Physics & Astronomy Department, University of Western Ontario, London ON, Canada; University of Ontario Institute of Technology, Faculty of Social Science and Humanities, Oshawa ON Canada

## Abstract

**Objectives:** Inconsistencies in the literature make it difficult to outline the relationship between exercise and cognition in young adults. Our aim is to better understand the relationship between prolonged aerobic and resistance exercise and cognitive abilities in sedentary young adults, and how this relationship is mediated by changes in cardiovascular fitness.

**Methods:** Twenty-three volunteers were recruited and assigned to two groups to complete one hour of continuous daily workout sessions of aerobic (SPIN) and anaerobic (SCSW) exercises over a 30 day period. Each subject was provided with a Polar-10 wearable to record the heart rate (HR) activity during the workout sessions. The workout sessions were completed during five consecutive days over four consecutive weeks. Each week, HR data were collected from the last workout session. Volunteers also completed a neurocognitive test battery (Cambridge Brain Sciences, CBS) each exercise session, including an additional baseline measure before exercise regime began.

**Results:** We found that memory, reasoning and verbal abilities improved throughout the aerobic, but not the resistance exercise program. We found a positive correlation between heart rate index (HRI) and memory and reasoning test scores. We also found a negative correlation between reasoning ability and HRM (heart rate mean), and heart rate skewness (SKW). The results of a regression model to predict memory and reasoning abiltiies revealed that memory was best predicted by HRI and HRM, while the reasoning ability was best predicted by only HRI.

**Conclusion:** Regular aerobic exercises improved specific cognitive performance and it was possible to predict the performance by employing the HR parameters HRI and HRM.

## Introduction

The health benefits of physical activity are well documented. Regular exercise is important for sustaining a healthy body-weight, and can help in the maintenance of normal blood lipid levels and blood pressure [1]. However, the benefits of exercise extend beyond physiological metrics; exercise has been shown to benefit mental health, improve self-esteem and self-confidence, and reduce symptoms of depression [2]. The benefits of exercise are so systematic that it is often recommended to prevent and manage various medical conditions, including cardiovascular disease, metabolic syndrome, osteoporosis, a number of neoplastic diseases, and various mental illnesses [1].

There is accumulating evidence that exercise can have drastic effects on various cognitive abilities, including improving learning and memory, delaying onset and magnitude of cognitive decline associated with various neurodegenerative diseases, and better outcomes associated with brain injury [3].

While the link between cognition and different types of exercise has been investigated in different age groups across the lifespan, most of this work focuses on the effects of aerobic exercise on cognition in older adults. In this cohort, there is evidence to suggest that older adults with greater cardiovascular fitness levels tend to experience less severe age-related decline in cognitive function [4], and in some cases, improvements to various aspects of cognition. For instance, aerobic exercise has been shown to benefit performance on tasks measuring attention, spatial memory, executive function, and processing speed in older adults [5].

Improvements in cognitive functioning were also accompanied by changes to the brain. Better cardiovascular fitness is also associated with reduced age-related neural degeneration of the prefrontal, parietal and temporal cortices and increased hippocampal volume[6]. These neural changes are accompanied by superior performance on tasks measuring memory [7], executive function, attention [8], and global intelligence [9].

Fewer studies have investigated whether exercise may lead to cognitive improvements in older adolescents and young adults. The relatively limited research focusing on this developmental period is surprising for two reasons: 1) approximately three-quarters of adolescents and young adults are insufficiently physically active [10], and over 80% of adolescents and young adults do not adhere to the recommended physical activity guidelines [11], and 2) the lack of physical activity can be major threat to cognitive health, potentially affecting various aspects of cognition, such as executive functioning, which continues to develop into adulthood [12]. Indeed, not all forms of exercise are beneficial to cognition[13].

Inconsistencies in the literature make it difficult to draw conclusions about the relationship between exercise and cognition in older adolescents and young adults. One reason is the limited overlap in the design parameters of the exercise interventions between studies. For instance, studies examining the effects of exercise on cognition differ in the type of exercise (i.e., aerobic, strength training and multimodal forms of exercise) and the duration of the exercise (i.e., acute and long-term) [6,14]. Another important factor is the considerable heterogeneity in the populations studied and the evaluated cognitive abilities. That is, most of the research in this area is confined to older adults, using a single type of exercise (e.g., aerobic training), and a limited set of cognitive tasks (e.g., memory tasks).

The current study aims to address these issues by: 1) focusing on older adolescents and young adults, 2) employing both aerobic and resistance-based exercise regimes in a longitudinal design (approximately over four weeks), 3) tracking physiological changes associated with each exercise regime using heart-rate variability, and 4) using Cambridge Brain Sciences (CBS) platform to assess cognitive performance. CBS is a set of 12 online tasks used to measure various aspects of cognitive function that collectively assess three cognitive domains: short-term memory (STM), reasoning and verbal abilities. Performance on the three cognitive domains have been validated based on data collected from tens of thousands of individuals. Neuroimaging data has demonstratee that each domain is supported by a separate brain networks [15]. Using the cognitive domains extracted from the CBS battery is ideal because the influence of different types of prolonged physical activity can be examined on three distinct and global aspects of cognition.

Studying young adults is fundamental to understanding the relationship between cognitive function and exercise because of the malleable nature and continuing development of cognitive functioning [12]. By eliminating the confounds of developmental and aging-related factors that affect cognition while using a comprehensive task battery, we set out to complete a more exhaustive evaluation of the effects on cognitive performance after introducing an exercise regime. Specifically, we aimed to delineate aspects of cognition that are most influenced by long-term aerobic and resistance exercise regimes in young adults. We hypothesized that participants in the aerobic exercise group would improve more on tasks relying on executive functions, such as working memory, while tasks relying on verbal and memory abilities would show the most improvement after completing the resistance-based exercise regime.

## Materials and methods

### Participants

Twenty-three volunteers were recruited in this study. The inclusion criteria required the volunteers to have a sedentary lifestyle. The participants were randomly assigned to two different groups: SPIN (1 males, age 22; 11 females, age 23 ± 3) and SCSW (5 females, age 23 ± 3). Participants (six in total) who did not complete a minimum of four exercise sessions and all cognitive testing sessions were removed;The SPIN and SCSW groups’ subjects were instructed to perform one hour of continuous daily workout sessions of aerobic and anaerobic exercises, respectively. Each subject was provided with a Polar 10 wearable to record the heart rate activity during the workout sessions, and scheduled the daily workout sessions to not interfere with personal daily activity. The workout sessions were performed for five consecutive days (from Monday to Friday) for four consecutive weeks (figure 1). For each week, HR data were collected from the last workout session.

**Figure 1.**
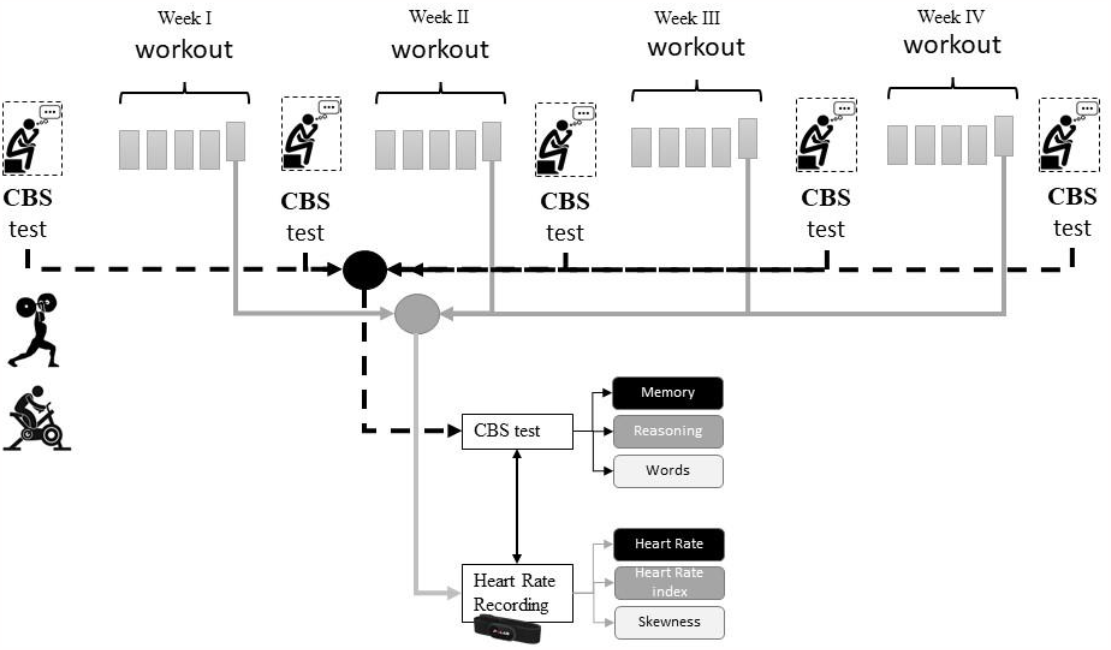
Scheme of the study. The volunteers were divided into SPIN and SCSW groups, corresponding to aerobic and anaerobic exercises, respectively. Each subject completed five workout sessions for week, for four consecutive weeks. HR parameters were collected from the last workout session of each week. The CBS tests were performed before starting with the workout sessions and after each weekly workout session. The two groups’ performance in CBS cognitive test were studied for the different domains (memory, reasoning, and words) and HR parameters. The HR parameters were collected during the last workout session of the week.

Five neurocognitive test batteries (Cambridge Brain Sciences, CBS) sessions were also performed; the first CBS test (CBS_0_) before starting with the workout sessions, and the others (CBS_1_-CBS_4_) the successive day following the last weekly workout session (figure 1).

### Neurocognitive assessment

CBS is a web-based neurocognitive assessment battery consisting of 12 tests that tap into a broad range of cognitive processes. The battery test is divided into three domains: reasoning skills, short-term memory, and verbal ability. The reasoning skill consists of 5 tests, the short-term memory of 4 tests, and the verbal ability of 3 tests, that include Feature Match, Odd One Out, Polygons, Rotations, Spatial Planning, Monkey Ladder, Paired Associates, Spatial Span, Spatial Search, Digit Span, Double Trouble, and Grammatical reasoning. CBS can be self-administered by individuals without the need for a neuropsychologist or other trained personnel. Each domain score is computed by the results of the related tests.

### HR recording

The heart rate (HR) signal was acquired by Polar H10 chest strap (www.polar.com/au-en, https://www.polar.com/en/science/whitepapers/h10-heart-rate-sensor-white-paper). The device collects the signal at 1 Hz with a sample tempo HR of 0.1 Hz (figure 2), providing time-stamped HR data based on the Network Time Protocol [16].

**Figure 2:**
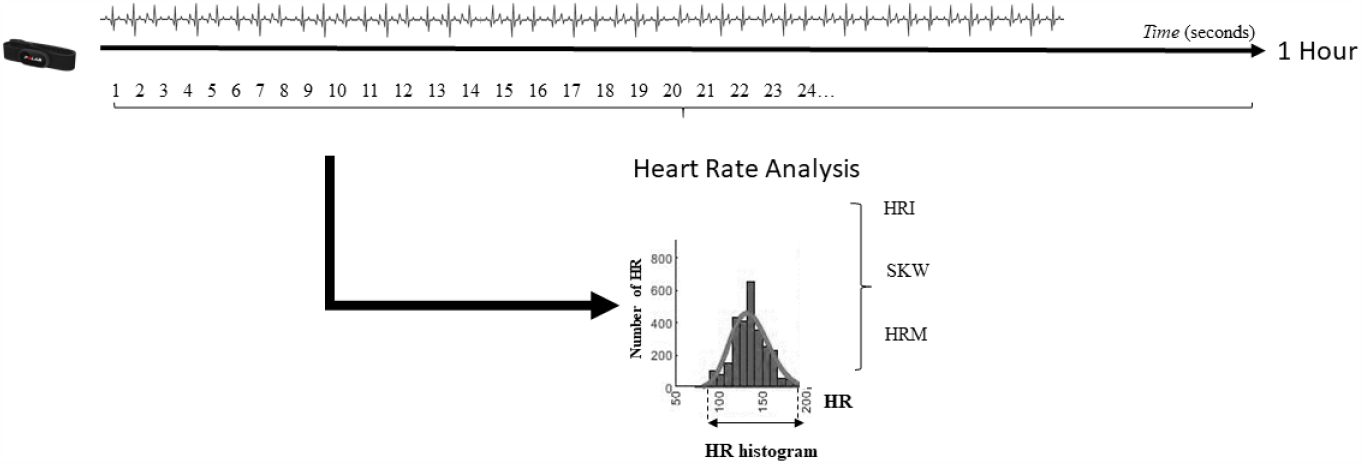
Scheme of the polar 10 acquisition data. Every second, the device provides the HR. From the collected data, HR triangular index (HRI), skewness (SKW), and HR mean (HRM) were extracted.

The HR collected data were controlled for missing outlier values. Missing or outlier values were corrected by the interpolation method.

HR data were analyzed by geometrical methods that represent an alternative to less easily obtainable statistical parameters for recording with a duration of at least 20 minutes and because less affected by the quality of recorded data.

For each workout session, HR mean (HRM), HR triangular index (HRI) and its skewness fitting curve (SKW) were calculated. The HRI was calculated as the integral of the density distribution (i.e., the number of all peak-to-peak (RR) intervals of the ECG) divided by the maximum of the density distribution. As the Polar H10 provides the HR every second, the HRI was calculated as the integral of the density distribution of the HR histogram divided by the maximum density distribution (figure 1).

## Statistical Analysis

For both groups, the first CBS test session of the three CBS domains was compared with the successive by Wilcoxon’s exact test. Moreover, for each session, the groups were compared for the different domains by Mann-Whitney exact test.

Similarly, for both groups, HRM, SKW, and HRI of the first workout session were compared with the successive by Wilcoxon’s exact test. The effect size r was calculated as the absolute value of Z/√(N) where Z is the Z-statistic of the statistical test, and N is the total number of subjects. The effect size results were considered: r<0.1 not significant; 0.1≤r<0.3 low; 0.3≤r<0.5 medium; r>0.5 high. Based on Bonferroni correction, the significance level for multiple comparisons was set at p<0.006. Correlations within HR parameters and between HR parameters and results in the different CBS cognitive domains were observed by Pearson’s correlation test. Linear regression analysis was applied to observe which HR variable is significant in predicting the CBS tests’ results in the different cognitive domains.

## Results

Six participants of the SCSW group were excluded from the analysis because of bad HR recording data due to incorrect device use or not completing the workout sessions.

### CBS sessions

No significant difference was found between SPIN and SCSW exercise groups for any of the scores in the three domains across all CBS sessions (CBS_0_-CBS_5_) (−2.0≤Z≤-0.3; 0.01≤*p*≤0.4; after Bonferroni Correction). However, we did find a significant difference between CBS_0_ and CBS_3_ in the memory (Z=-2.6, p=0.003, r=0.53) and reasoning domain (Z=-2.5, p=0.003, r=0.51) for the SPIN group. The SPIN group also showed a significant difference between CBS_0_ and CBS_2_ (z=-2.5, p=0.005, r=0.51) and CBS_0_ and CBS_4_ for the verbal domain (z=-3.1, p<0.0001, r=0.53). All comparisons were based on the Wilcoxon’s exact test (figure 3).

**Figure 3:**
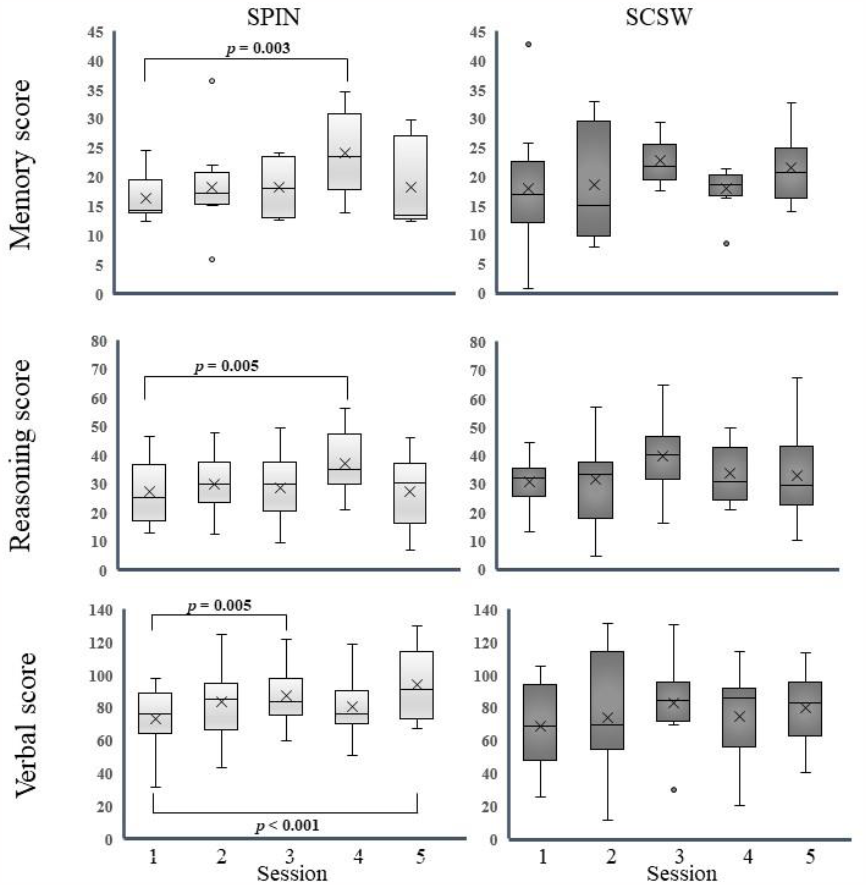
Boxplot of CBS test. SPIN: aerobic workout, SCSW anaerobic workout. Into the boxplot: the median (horizontal line), the mean (x), and the number of the CBS session (1-5). Session 1 is the CBS test before the workout session.

**Figure 4:**
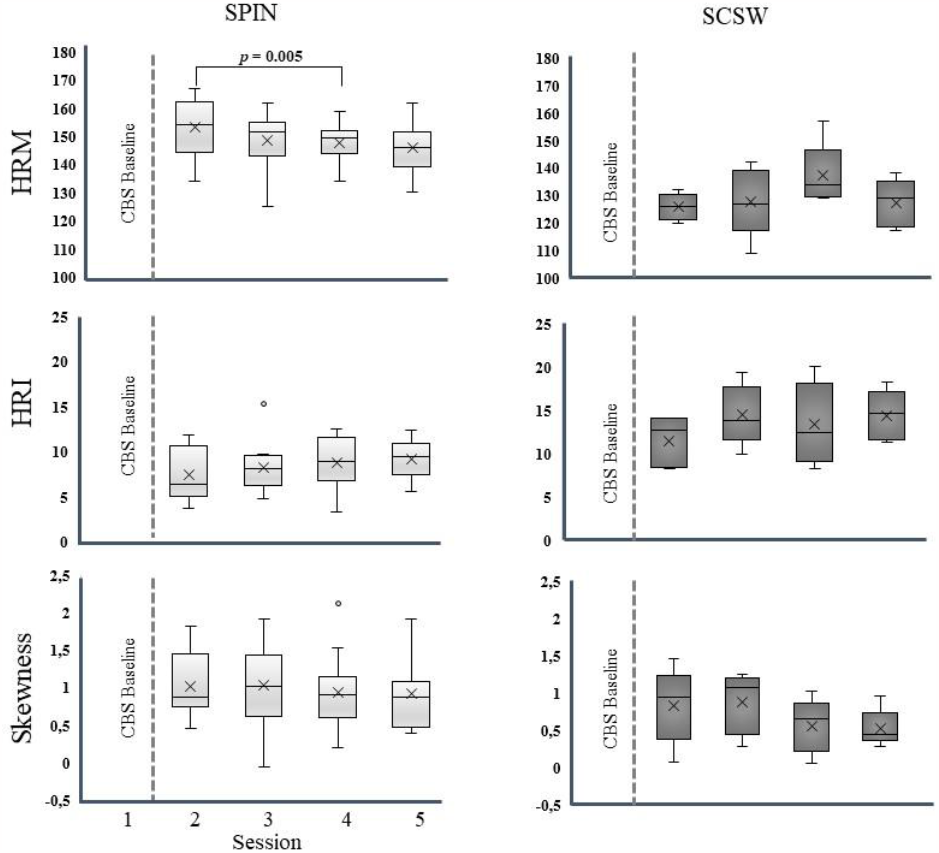
Boxplot of HR measures. SPIN: aerobic workout, SCSW anaerobic workout.HRM: heart rate mean, HRI: heart rate triangular index, SKW: skewness fitting curve of HRI. Into the boxplot: the median (horizontal line), the mean (x), and the number of the CBS session (1-5). Session 1 is the CBS test before the workout session with no HR recording.

### HR measures

Comparing HRI, HRM, and SKW between the first workout sessions with the following three sessions, we found a significant difference at Wilcoxon’s exact test between HRM_1_ and HRM_3_ (Z=- 2.5, p=0.005, r=0.43) for the SPIN group. No differences were found in the other conditions for SPIN (−2.1≤Z≤-0.8, 0.01≤p≤0.4) and SCSW (−2.0≤Z≤-0.1, 0.03≤p≤0.5) groups.

Comparing SPIN and SCSW for HRI, HRM, and SKW, at Mann-Whitney exact test, significantly higher values in SCSW group were found for HRI_1_ (Z=-2.8, p=0.001, r=0.49) and HRI_4_ (Z=-2.8, p=0.001, r=0.49), and in SPIN groups for HRM_1_ (Z=-3.1, p=0.0001, r=0.55) HRM_2_ (Z=-2.6, p=0.004, r=0.45) and HRM_4_ (Z=-2.8, p=0.001, r=0.48). No significant differences were found for the SKW (−1.7≤Z≤-0.3, 0.05≤p≤0.4).

At the Spearman’s correlation test, an inverse correlation was found between HRI and SKW (Rho=- 0.65, p<0.0001), and between HRI and HRM (Rho=-0.64, p<0.0001), while a positive correlation was found between SKW and HRM (Rho=0.34, p=0.005) (figure 5).

**Figure 5:**
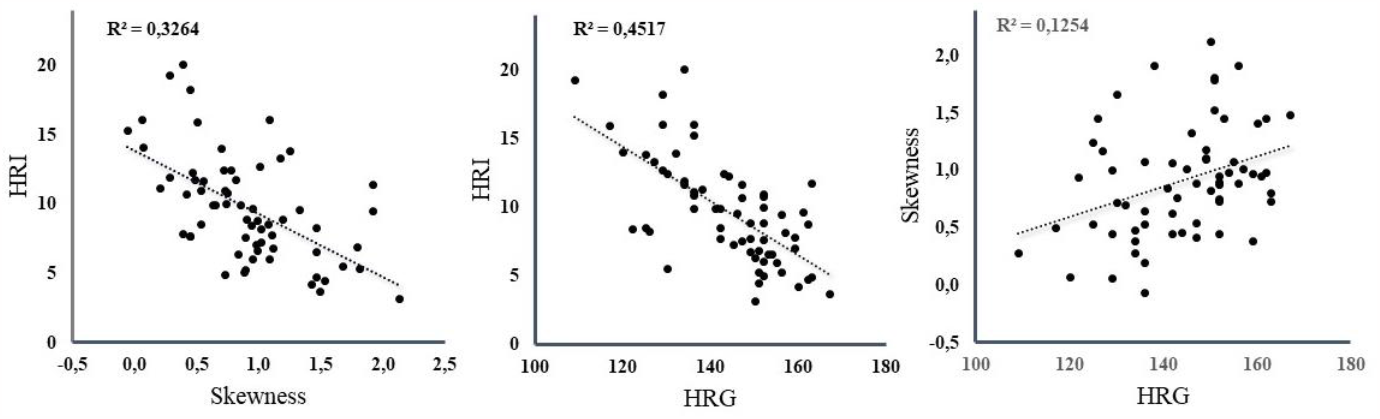
scatterplot of the HRI, SKW, and HRM of the workout sessions for all subjects. The SKW ranged from negative to positive values and is dependent on direction: positive SKW has negative values, negative SKW has positive values.

### Correlations between CBS tests and HR measures

A significant positive correlation was found between HRI and the scores in memory and reasoning tests (Spearman’s Correlation: Rho=0.33, p=0.004, and Rho=0.37, p=0.001, respectively). A negative correlation between the scores in the reasoning tests and HRM (Spearman’s Correlation: Rho= -0.36, p=0.002) and SKW (Spearman’s Correlation: Rho=-0.35, p=0.003) was also found significant (figure 6).

**Figure 6:**
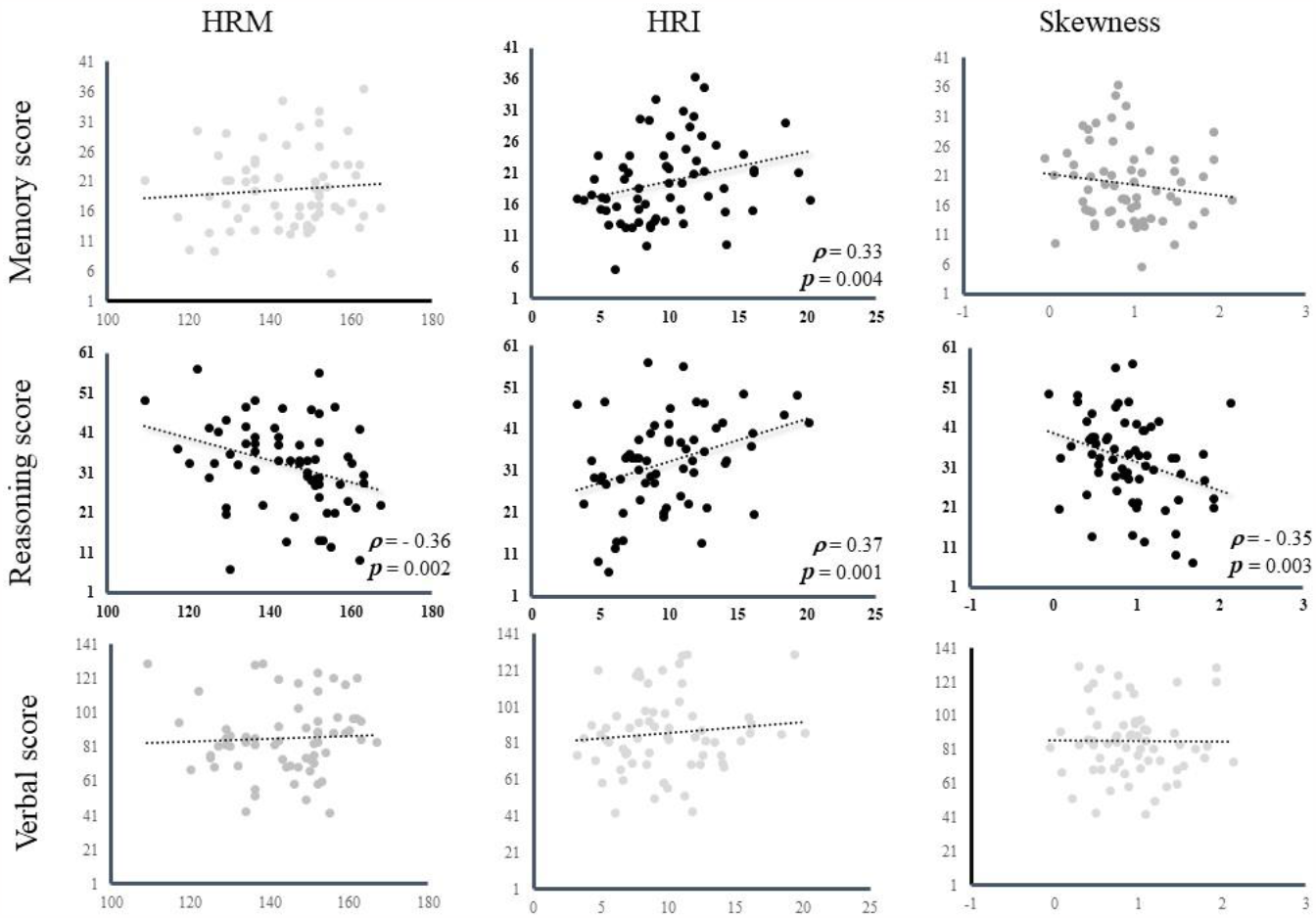
scatterplot of the scores in the different cognitive domains after the workout sessions with the SKW (right column) and HRI (central column) and HRM (left column) for all subjects. In bold the significant correlations with p-value and Rho’s Spearman.

**Figure 7:**
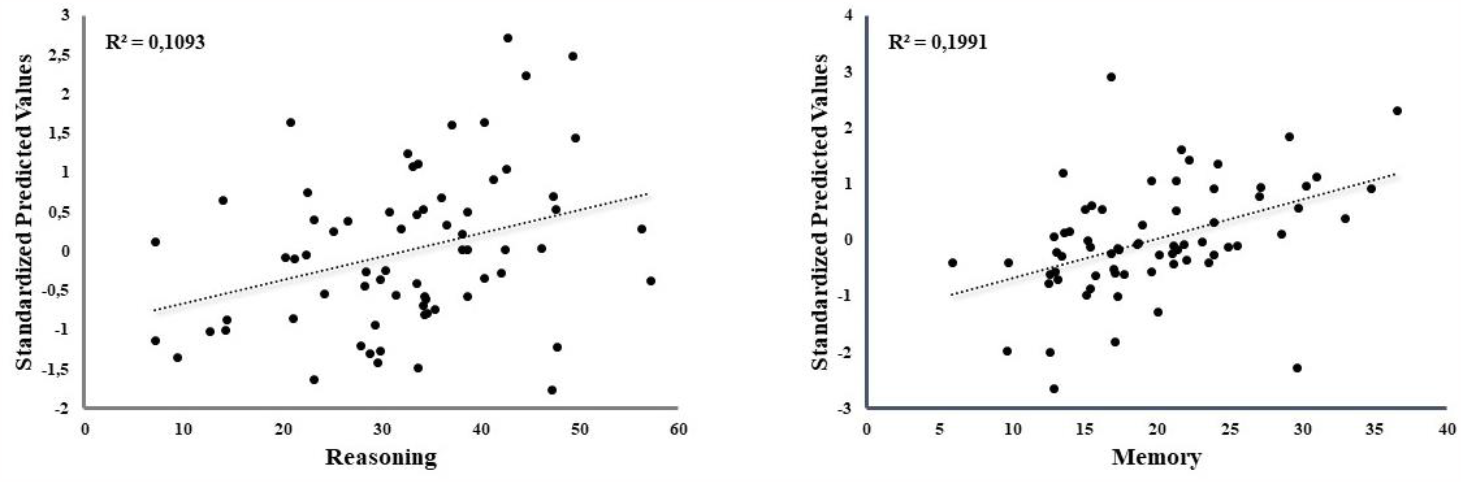
scatter plot of the linear regression. In the x-axes, the scores in the cognitive tests, and in the y-axes the standardized predicted values of the linear regression model. Based on the HR parameters, the predicted score in the reasoning domain is equal to 22.981 +0.968(HRI), and predicted score in the memory domain is equal to -24.111 +1.017(HRI)+0.236(HRM).

### HR measures predict cognitive results

In the multiple linear regression analysis, HRI, HRM, SKW, and workout variables were used to predict memory, reasoning, and verbal results in the cognitive tests. No significant regression model was found for the verbal domain. A significant regression model was instead recognized for the memory (F=2,65=8.078, p=0.001, R^2^=0.20) and reasoning domain (F=1,66=8.102; p=0.001; R^2^=0.11). The prediction of the score in the memory domain was based on the HRI (p<0.0001) and HRM (p=0.002), with the predicted score equal to -24.1 + 1.0*HRI + 0.2*HRM, whereas the prediction of the score in the reasoning domain was based only on the HRI (p=0.006), with the predicted score equal to 23.0 + 1.0*HRI.

## Discussion

Prolonged aerobic exercise (i.e., spin class) was associated with significant improvements to memory, reasoning and verbal ability, with the improvements to these cognitive abilities emerging as soon as three weeks of regularly attending spin classes. In addition to exercise improvements to cognition, prolonged exercise was related to changes in heart-rate; mean heart-rate was significantly lower after three weeks of aerobic exercise relative to mean heart-rate at the start of the exercise regime. Importantly, reduced mean heart-rate was correlated with reasoning abilties, that is, improved reasoning abilities over time was associated with improved cardiovascular health. Morevoer, subcomponents of heart-rate data were also associated with changes to cognitive functioning; the heart-rate triangular index, and the skewness fitting curve, both markers of sympathovalgal interaction [17], were associated with improvements to reasoning and memory abilties throughout the exercise regime.

Our results are consistent with previous work demonstrating improved cognitive functioning following aerobic exercise [18]. For instance, Miles and colleagues [19] found a strong relationship between increasing HR during aerobic exercise and performance on memory tasks. In particular, Miles and colleagues observed that a change in HR between learning and retrieval following the rest-exercise condition (or vice versa) can have a powerful effect on improving the memory test. The increased heart rate during exercise could be associated with a more intense physical effort with a consequent improvement in the modulation of cardiac activity and in the ANS response [20], explaining the role of HRM in predicting the performance for the memory test.

One advantage of the approach of examining subcomponents of the heart-rate is that we were able to isolate which aspects of cognition are most strongly linked to different metrics of cardiovascular functioning. Kouidi and colleagues [17] showed the effects of the athletic training on the HRI and its positive correlation with the vagal activity. That we found HRI had a positive relationship with reasoning and memory indicates that increased vagal control is linked to better performance in these cognitive field, as suggested in other studies [21,22], but not to verbal abilities [23]. At the same time, we found HRM and SKW negatively associated with reasoning abilities; lower values for HRM and a more normal distribution of the HR (smaller values for SKW) were linked to better performances on tasks that measure reasoning. This suggests a probabily better breathing modality, that influences the vagal control in reasoning performances [24].

Beyond identifying relationships between cognition and various metrics of HR, we also identified weighted contributions of HR metrics to predict cognition. Reasoning ability was best predicted by the HRI only, while memory was best predicted by a combination of HRI and HRM. This would suggest a different autonomic modulation linked to a more stress-control during the cognitive memory test [25].

Our results complements previous findings that aerobic exercise has been shown to significantly improve various aspects of cognition [26]. For example, six months of aerobic resulted in improvemnts to learning and memory in older healthy male and female participants [27]. These benefits also extend to clinical populations; intensive aerobic exercises improved processing speed and divided attention in post-stroke patients with cognitive impairments [28], and patients with mild cognitive impairment [29]. However, our results suggest that the same benefits to cognitive functioning apply to young sedentary adults.

What are the potential mechanisms underlying our results demonstrating improvements to cognition and cardiovascular health after prolonged aerobic exercise? One possibility is that the reduction in mean HR, an increase in HRI and a positive shift of the HR skewness during the exercise regime suggests increased vagal modulation of the ANS, which is an adaptive response to physical stress [30]. The adaptive response of the ANS is part of a more complex network. This network, known as Central Autonomic Network (a model that describes the brain-heart two-way interaction), is also involved in the behavioral response [31]. Another potential mechanism is that aerobic exercise may have triggered protective brain functions [3]; for example, a neurotrophic factor involved in the mediation of the beneficial effects of exercise on brain plasticity is the brain-derived neurotrophic factor (BDNF), a molecule that belongs to a family of neurotrophins [3]. BDNF has a central role in survival and differentiation of neuronal populations during development [32] and is involved in plastic changes associated with improvements to learning and memory [33].

Despite finding strong links between cardiovascular and cognitive changes during prolonged aerobic exercise, we did not find the same pattern of results associated with a strength-based exercise regime. That is, completing four weeks of sculpt and sweat was not associated with significant improvements to any aspect of cognition, or was it associated with changes to any of the measures of cardiovascular health. One reason for this difference may be because of the downstream effects each type of exercise has on the Cardiovascular Nervous System (CNS) and ANS. That is, exercise is defined as any activity that uses large muscle groups, can be maintained continuously, and is rhythmic in nature [34]. Anaerobic exercise is characterized by an intense physical activity of brief duration, fueled by the energy sources within the contracting muscles and independent of the use of inhaled oxygen as an energy source [34]. During aerobic exercises, the cardiorespiratory system supplies oxygen that the skeletal muscle system utilizes. In contrast, during resistance exercises, without the use of oxygen, the cells revert to the formation of ATP via glycolysis and fermentation. Physical exercises improve hemodynamic, hormonal, metabolic, neurological, and respiratory function [1]. Regular aerobic training seems to lead to an improvement of vagal modulation, observable with a reduction of the heart rate and increasing of the HRV at rest as well as during submaximal exercise [35]. In general, aerobic exercises increase parasympathetic activity, while anaerobic exercises stabilize the function of the ANS [36].

Several studies reported the different effect of aerobic and anaerobic activity on cognitive performances evidencing that the first improve the performance in attention and memory tasks [37,38] while the second in memory, verbal concept formation, and selective attention [39,40].

Moreover the effects of esercise on the cognitive functions seem to be dependent on the amount of time dedicated to the fisical training. In that regard, it was found that 20 minutes of aerobic exercise increase reaction time and cognitive flexibility (17-20), while 30 minutes of resistance exercise improve performance in executive function (15-16).

Meta-analysis works of McMorris and colleagues [41,42] report that moderate intensity of excercise could increase peripheral catecholamine concentrations inducing vagal/nucleus tractus solitarius pathway activation with a positive effect on reaction time and cognition.

These cognitive functions involve planning, scheduling, updating, and initiation ability, processed mainly in the prefrontal cortex [22,43]. Aerobic exercises, increasing the heart rate level, increase arousal level and activation of cortical areas involved in cognitive performances [44,45]. Resistance exercises are also correlated with cognitive improvement but with a different mechanism, where the increase in heart rate level is linked to increased hormone concentration[46].

Comparing aerobic versus resistance exercises, if Alves and colleagues [47] found that both improve in middle-age women speed processing and inhibition control in the Stroop test, Pontifex and Hillman [22] found that improvement of speed processing of a working memory task improves after aerobic exercises but not resistance exercises. Similarly, Dunsky and colleagues [48] found a positive effect on various tasks involving attention due to the acute aerobic, but no resistance, exercise.

A review by Netz [49] suggests that physical training 1) affects cognition via the improvement of cardiovascular fitness, 2) affects neuroplasticity and cognition globally, and 3) that the intensity of training enhances neuroplasticity improving cognition. Differently, motor training seems 1) to affect cognition directly, 2) that the increasing brain neuroplasticity in affecting cognition is task-specific, and 3) that the relationship between exercise and cognition is affected by motor complexity.

Our findings highlight that adaptation to the ANS in response to aerobic exercise is associated with improved performance in memory and reasoning abilities supporting the different effects of aerobic and resistance exercises. Unique to our study, this relationship was assessed with heart-rate recording during the physical effort, in contrast to other studies based on the heart rate variability analysis (i.e., fluctuation in the time intervals between adjacent heartbeats). Despite not obtaining RR intervals (the peaks between two consecutive R-wave of the QRS complex of the ECG measured in milliseconds), we showed that the HRI, HRM, and SKW could be used equally well to quantify ANS response changes. Importantly, these measures are less affected by movement artefacts that, during physical exercise, depending on the intensity and chosen exercises [50], can highly perturb RR interval acquisition and introduce biases in the HRV signal.

This study contains important limitations. First, our sample was made up of majority female participants; in fact every participant except one was female. Until a sample of male participants are tested in a similar design, our results can only be generalized to older adolescent/young adult females. Second, attrition rates for longitudinal exercise studies are hight, and this was no exception for our study; as a result only 5 participants in the strength-based exercise group (SCSW) produced usable data. The sample size of the SCSW group could represent the reason we failed to find improvements in cognition following anaerobic training. Future studies with more power are necessary for replicating our results.

